# Wheat grain width: A clue for re-exploring visual indicators of grain weight

**DOI:** 10.1101/2021.10.13.464205

**Authors:** Abbas Haghshenas, Yahya Emam, Saeid Jafarizadeh

## Abstract

Mean grain weight (MGW) is among the most frequently measured parameters in wheat breeding and physiology. Although in the recent decades, various wheat grain analyses (e.g. counting, and determining the size, color, or shape features) have been facilitated thanks to the automated image processing systems, MGW estimations has been limited to using few number of image-derived indices; i.e. mainly the linear or power models developed based on the projected area (*Area*). Following a preliminary observation which indicated the potential of grain width in improving the predictions, the present study was conducted to explore potentially more efficient indices for increasing the precision of image-based MGW estimations. For this purpose, an image archive of the grains was processed, which was harvested from a two-year field experiment carried out with 3 replicates under two irrigation conditions and included 15 cultivar mixture treatments (so the archive was consisted of 180 images taken from an overall number of more than 72000 grains). It was observed that among the more than 30 evaluated indices of grain size and shape, indicators of grain width (i.e. *Minor* & *MinFeret*) along with 8 other empirical indices had a higher correlation with MGW, compared with *Area*. The most precise MGW predictions were obtained using the *Area*×*Circularity*, *Perimeter*×*Circularity*, and *Area/Perimeter* indices. In general, two main common factors were detected in the structure of the major indices, i.e. either grain width or the *Area/Perimeter* ratio. Moreover, comparative efficiency of the superior indices almost remained stable across the 4 environmental conditions. Eventually, using the selected indices, ten simple linear models were developed and validated for MGW prediction, which indicated a relatively higher precision than the current *Area*-based models. The considerable effect of enhancing image resolution on the precision of the models has been also evidenced. It is expected that the findings of the present study improve the precision of the image-based MGW estimations, and consequently facilitate wheat breeding and physiological assessments.

## 1. Introduction

Although number of grains per unit of area is known to be the most important component of wheat yield (García et al., 2014; and Slafer et al., 2014), grain weight and its related features (e.g. size and shape) are still under consideration of the researchers for improving the yield capacity (e.g. see Ramya et al., 2010; Williams et al., 2013; Brinton et al., 2017; and Alemu et al., 2020). Accordingly, wheat grain has been well-explored visually in the last decades, either using uncomplicated methods and 2D indices (Braadbaart & van Bergen., 2005; Firatligil-Durmuş et al., 2010; Gegas et al., 2010; Zapotoczny, 2011; Williams et al., 2013; and Whan et al., 2014) or employing more complex techniques of 3D reconstruction (Mabille & Abecassis, 2003; Strange et al., 2015; and Le et al., 2019). The techniques utilized for this purpose can be categorized under the term of high-throughput phenotyping (HTP), which has been emerged as an efficient paradigm in response to the need for keeping the feasibility of investigations in the current complex and large-scale breeding programs.

The most frequent sensors used in HTP are the efficient, inexpensive, and widely available RGB cameras (Araus et al., 2018). A simple processing of an RGB image of grains along with utilizing appropriate indices of size, color, and shape, can thoroughly and rapidly quantify the phenotype of grain samples. It seems most reasonable to select the projected area (*Area*) as the most relevant image-derived index for estimating grain weight; as this indicator provides a 2D representation of the 3D grain size (compared with the one-dimensional criteria e.g. grain width or length). Accordingly, studying the relationship between the area and weight of individual grains, Kim et al. (2021) introduced a single power model equation for estimating wheat grain weight, (i.e. *weight* = *area*^1.32^), which provided a higher precision compared with the linear model.

In a preliminary analysis conducted with the aim of evaluating the variations of grain size and shape in wheat cultivar mixtures, it was observed accidentally that grain weight had a relatively higher correlation with grain width, compared with the well-assessed index of projected grain area. This observation encouraged a more comprehensive analysis for potentially improving the image-based estimation of wheat grain weight. Therefore, the purpose of the present study was (i) assessing and documenting the relative advantage of grain width, and also (ii) seeking more efficient image-derived indices for predicting grain weight.

## 2. Materials and Methods

### 2.1. Field experiment

The images of wheat grains were selected from an archive of a two-year field experiment conducted with the aim of studying the responses of cultivar mixtures with various ripening patterns to normal and post-anthesis water stress conditions (see Haghshenas et al., 2021). The experiment was conducted during 2014-15 and 2015-16 growing seasons at the research field of the School of Agriculture, Shiraz University, Iran (29°73′ N latitude and 52°59′ E longitude at an altitude of 1,810 masl). Mixture treatments were 15 mixing ratios of four early- to middle-ripening wheat cultivars (Chamran, Sirvan, Pishtaz, and Shiraz, respectively) including the 4 monocultures and their every 11 possible mixtures, which were grown with 3 replicates under two well-irrigation and post-anthesis deficit-irrigation conditions. The experimental design was RCBD (Randomized Complete Block Design) in which all the 90 (2×2 meter) plots were arranged in a lattice configuration with 1 meter distances. Plant density was 450 plants/m^2^ and seeds were mixed in each year with equal ratios (i.e. 1:1, 1:1:1, and 1:1:1:1 for the 2-, 3-, and 4-component blends, respectively), considering their 1000-grain weights and germination percentages. The planting date in the first and second growing seasons were November 20 and November 5, respectively; and based on the soil test, 150 kg nitrogen/ha was applied (as urea) in three equal splits i.e. at planting, early tillering, and anthesis. No pesticide was used and weeding was done by hand once at stem elongation.

Irrigation interval was 10 days based on local practices, and the amount of irrigation water was estimated using the Fao-56 Penman-Monteith model with local corrected coefficients which was reduced to 50% of evapo-transpirational demand from the first irrigation after anthesis. Late in the season, plants were harvested from the center of plots and yield components were estimated using a laboratory thresher and weighing.

### 2.2. Imaging

Images were taken from the archive of an exclusively designed laboratory system (Visual Grain Analyzer, VGA), which was equipped with a Logitech HD Pro Webcam C920 mounted on an adjustable arm, a glass table with a 60×60 cm flicker-free white LED panel beneath it as the light source, and a professional software written in C# for real-time screening of the grains. Imaging was carried out for other purposes, so the properties were not necessarily designed for the present study. Accordingly, images were taken under ambient light from 40 cm above the samples, and the image dimensions were 960×720 pixels (i.e. the original resolution was ≈ 7 MP). For each experimental plot, more than 400 grains were sampled randomly and arranged on the imaging table using a Vacuum Seed Counter, so that there was no contact between the grains. Therefore, the total dataset (including 90 images for each year) was consisted of the data of more than 72000 single grains. Immediately after imaging, the grains of each image were weighed using a A&D EK-610i (d=0.01 g) weighing balance. Mean grain weights were calculated by dividing the sample weight by the number of grains.

### 2.3. Image processing

Since the VGA system has not been commercialized or released yet, and also the analyses had to be kept reproducible, only the data of grain size (for conversion of pixel to mm) was taken from this system; and all of the image analyses were carried out using ImageJ version. 2.1.0/1.53c (Schindelin et al., 2012). First, the grains were segmented from the background using the *Color thresholding* tool (Image > Adjust > Color thresholding). The thresholding method and color space were set as “Default” and HSB, respectively. Thereafter, size and shape features of grains were calculated using the *Analyze particles* tool. For this purpose, the attended features were selected in the *Set Measurements* menu (*Analyze* > *Set Measurement*), and *Analyze Particles* was run. Before running, the “*Show Ellipses*” option was selected, and no size or circularity filtering was applied on the sample. The output tables were saved as .csv files and used for next analysis. As will be described later in the Results section, it was found out that enhancing the image resolution could improve the estimations. Therefore, in another analyses, before running the “*Analyze Particles*”, the resolution of images was enhanced using the Bicubic algorithm and by factor of 10 (i.e. both image dimensions were multiplied by 10, so the image resolution was increased 100 times). Resizing the images was carried out using the *Batch* processing tool (Process > Batch > Convert; and *interpolation* and *scale factor* were set to Bicubic & 10, respectively).

Using the output of image processing, the averaged values of basic features of size and shape were calculated for each image, and the correlation of these visual indices with MGW were evaluated. The examples of basic indices included *area*, *perimeter*, the major and minor axes of the best fitted ellipses to the grains (*Major* & *Minor*; also see Williams et al., 2013), minimum (*MinFeret*) and maximum (*Feret*) caliper diameter, *Circularity* (a value between 0 to 1 for an infinitely elongated shape to a perfect circle), *solidity* (the ratio of area to the convex hull area), etc. Besides the basic features, the correlation of MGW with several synthesized indices were also tested; which were the products or ratios of the basic indices. *A_1_* and *A_2_* were among the instances of synthesized indices which are the products of the 5 most efficient basic indices. The full list of the evaluated indices is represented in Table 1. Also for more detail of the definitions and formulae, see https://imagej.nih.gov/ij/docs/guide/146-30.html. Linear correlations of MGW with the visual indices were compared with those of the two control criteria i.e. *Area* and *Kim index* (*Area*^1.32^; taken from the paper of Kim et al., 2021), and the indices with a higher correlations than the controls were selected as the final indicators of MGW. Using each selected index, a linear model for prediction of MGW was developed and evaluated. Although the analyses were based on the number of pixels (as the unit of dimension), in order to generalize the model parameters, outputs were also converted into mm using the data of VGA system. Moreover, ten-fold cross-validation (K = 10) was used in Rapidminer (Version 9.9) to validate the results of datamining models, in which the default values and settings of the software were chosen. All other analyses, including correlating, Principal Component Analysis (PCA), and fitting the linear models were carried out using XLSTAT (Version 2016.02.28451; Addinsoft).

**Table 1.** List of the empirical image-derived indices tested in the present study.

| Preliminary indices | Selected indices |
| --- | --- |
| Area | Area |
| Perimeter (Perim.) | Minor |
| Major | MinFeret |
| Minor | Area/perim. |
| Circularity (Circ.) | Area×Circ. |
| Feret | Minor/Solidity |
| skewness (Skew) | MinF/Solidity |
| kurtosis (Kurt) | Area×Solodity |
| MinFeret (MinF) | Perimeter×Circ. |
| Aspect ratio (AR) | A1 (Area×Perim.×Circ.×Solidity×MinF) |
| Round | A2 (Area×Perim.×Circ.×Solidity×Minor) |
| Solidity | Kim index |
| Minor/Major |  |
| MinF/Feret |  |
| Area/MinF |  |
| Area/Minor |  |
| MinF/Minor |  |
| Area/prim. |  |
| Minor/Perim. |  |
| MinF/Perim. |  |
| Area/(Perim.^2) |  |
| MinF×Area/Perim. |  |
| Area/MinF |  |
| Area/Minor |  |
| Circ.×Solidity |  |
| Area×Circ. |  |
| MinF×Circ. |  |
| MinF/Solidity |  |
| Feret/Solidity |  |
| Area×Solodity |  |
| Feret×MinF×Solidity |  |
| Perimeter×Circ. |  |
| A1 (Area×Perim.×Circ.×Solidity×MinF) |  |
| A2 (Area×Perim.×Circ.×Solidity×Minor) |  |
| Kim index |  |

At the first step, the correlations between mean grain weight and the preliminary image-derived indices were tested. Then, the indices with a higher correlation coefficients (R) than those of the two control indices, i.e. "Area" and "Kim index", were selected for further analyses. Kim index (i.e. $Area^{1.32}$ ) was derived from the study of Kim et al., 2021. For definition of the other basic indices, see the ImageJ user guide on "Analyze particles..." at https://imagej.nih.gov/ij/docs/guide/146-30.html

## 3. Results

As shown in Fig. 1, enhancing the image resolution improved the quality of grain segmentation and ellipse fitting, considerably. This improvement was consequently reflected in the precision of the correlations and linear models developed for prediction of MGW (which will be discussed later).

**Figure 1.**
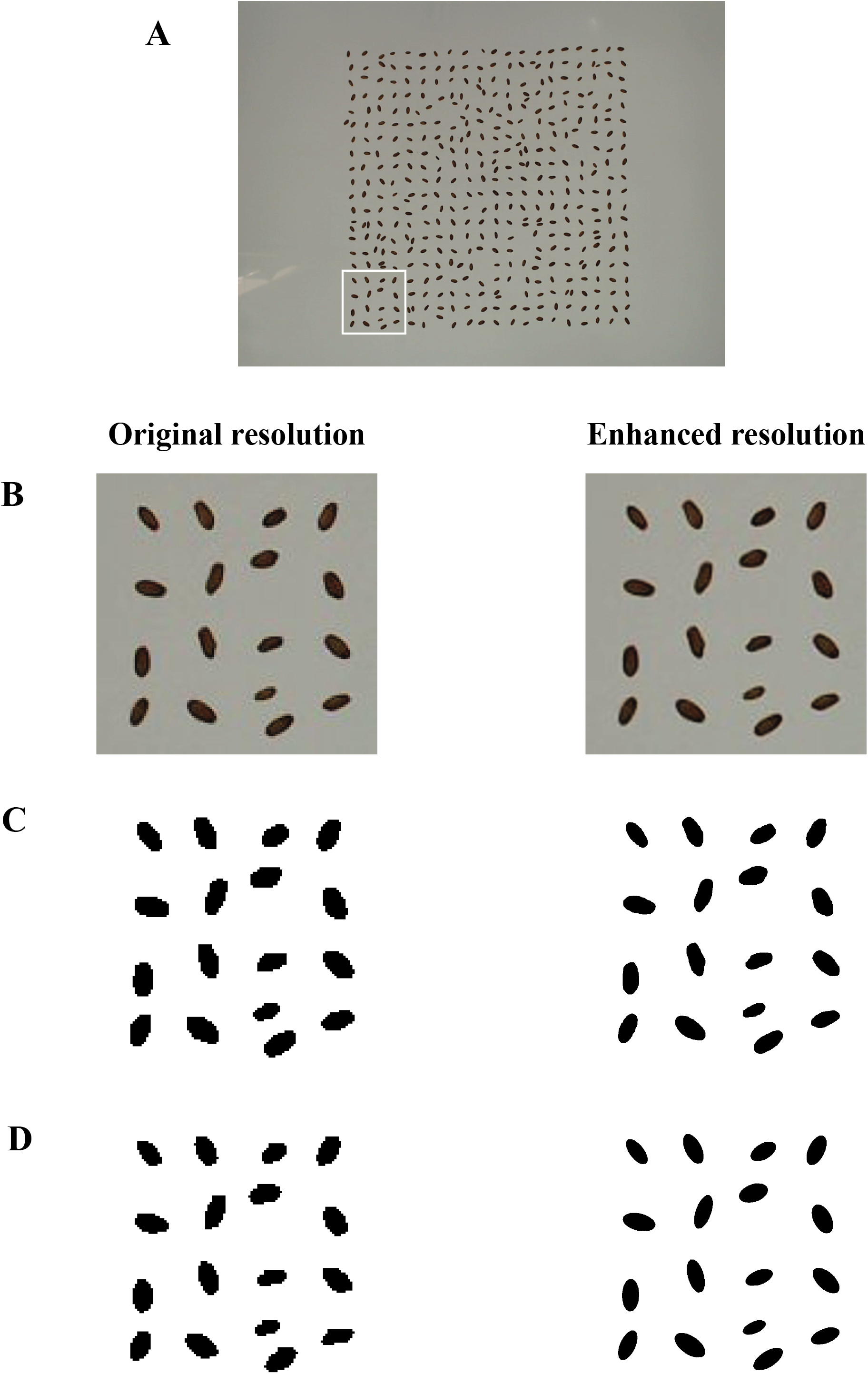
Output of image segmentation for extracting grains and fitting the best ellipses. (A) A single image from the archive with more than 400 wheat grains. As an example, the grains in the white frame are processed in the next parts of the figure. (B) Output of resolution enhancement; (C) Result of image segmentation. A same thresholding is used for both resolutions; (D) Fitting the best ellipses to the single grains.

Principal component analysis (Fig. 2) indicated that in comparison with area (R=0.905), the grain width had a stronger relationship with MGW; regardless of which width indicator was used (R=0.0921& R=0.916 in the cases of using *Minor* and *MinFeret*, respectively).

**Figure 2.**
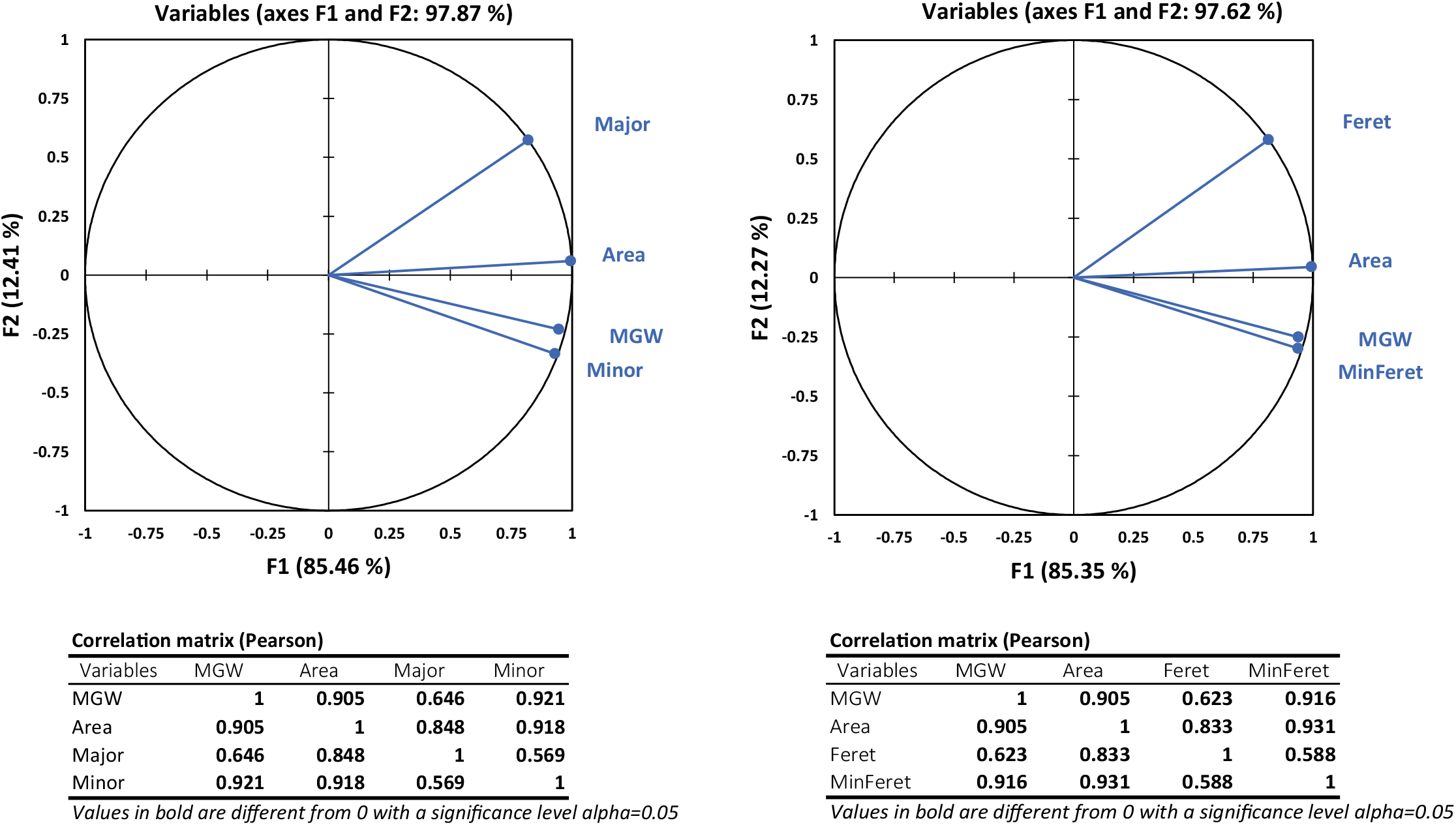
Principal Component Analysis (PCA) of mean grain weight (MGW) and basic image-derived indicators of grain size, i.e. major and Feret (indices of grain length), minor and minimum Feret (indicators of grain width), and area. Obviously, the one-dimensional indicators of grain width reflect the variations of MGW more precisely than the two-dimensional factor of area.

Besides the two control indices i.e. *Area* and *Kim index*, the correlation of MGW with 33 other preliminary indices were also tested; among which 10 indices with comparatively higher correlations than the two controls were selected for further analyses (Table 1). Figure 3 shows the correlations between MGW and the selected indices derived from the images with enhanced resolutions. The indices of *Area*×*Circ.*, *Perim.*×*Circ.*, and *Area/Perim.* had relatively stronger relationships with MGW. Table 2 also indicates the variations in the correlation coefficients (R) in various environmental conditions. It is obvious that almost in every conditions, the selected indices have a comparatively higher relationship with mean grain weight, compared with *Area* and *Kim index*. Again, the three indices mentioned before (i.e. *Area*×*Circ.*, *Perim.*×*Circ.*, and *Area/Perim*) had the highest R values, almost in every conditions. Moreover, in consistency with the fact shown in Fig. 1, the enhanced resolution improved the correlation considerably.

**Figure 3.**
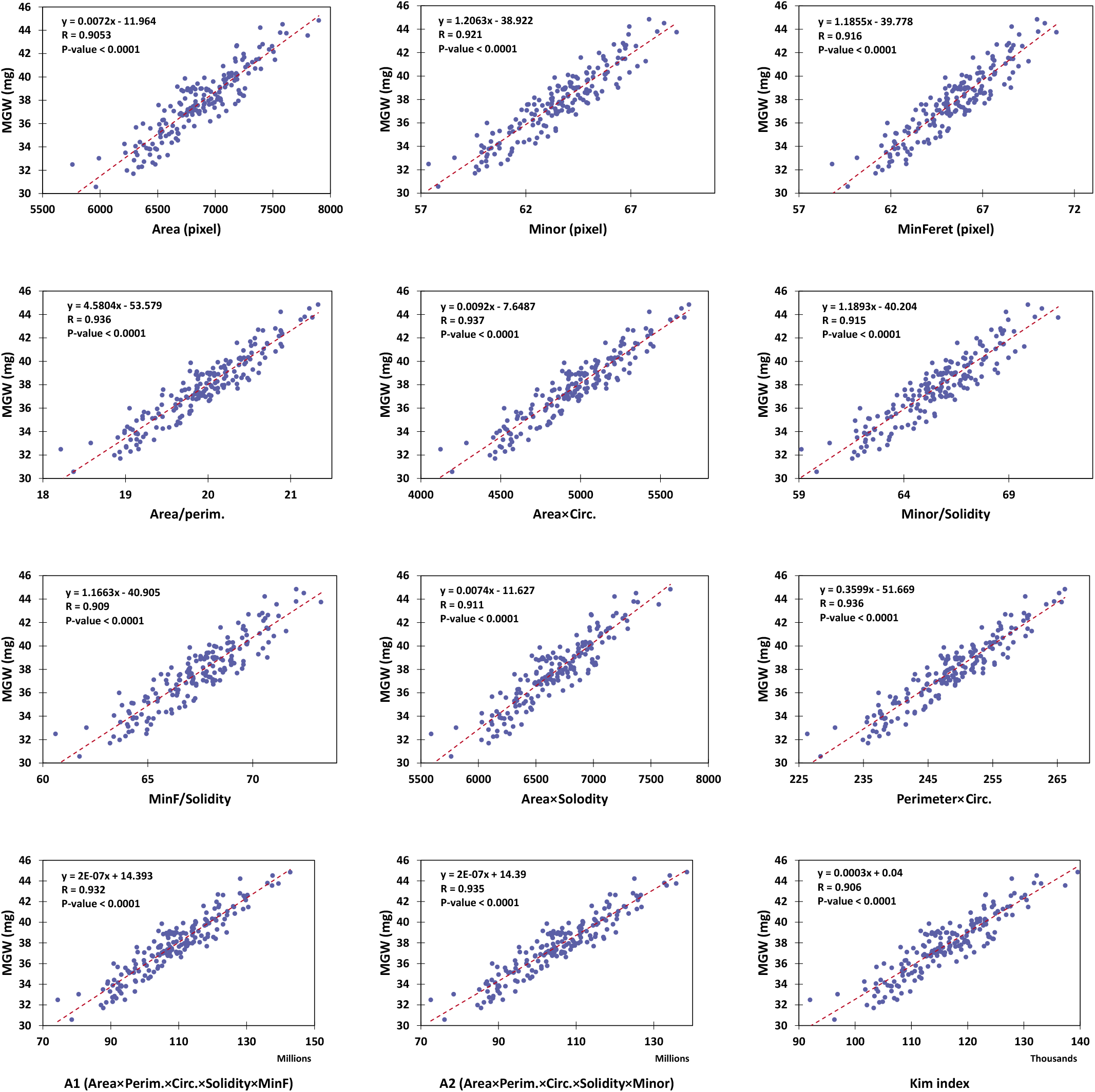
The correlations between mean grain weight (MGW) and image-derived indices. Here, the images with enhanced-resolution were used.

**Table 2.** The correlation coefficients (R) of mean grain weight (MGW) and image-derived indices.

| Resolution | Indices | Overall | 1<sup>st</sup> year | 2<sup>nd</sup> year | WI (two years) | DI (two years) | 1<sup>st</sup> Y. WI | 1<sup>st</sup> Y. DI | 2<sup>nd</sup> Y. WI | 2<sup>nd</sup> Y. DI |
| --- | --- | --- | --- | --- | --- | --- | --- | --- | --- | --- |
| Original resolution | Area | 0.8740 | 0.8382 | 0.8813 | 0.8092 | 0.8486 | 0.8117 | 0.7898 | 0.7805 | 0.8113 |
|  | Minor | 0.8790 | 0.9044 | 0.9034 | 0.7805 | 0.8549 | 0.8751 | 0.8621 | 0.7860 | 0.8310 |
|  | MinFerret | 0.8833 | 0.9030 | 0.9086 | 0.7930 | 0.8567 | 0.8664 | 0.8579 | 0.7897 | 0.8488 |
|  | Area/perim. | 0.8920 | 0.9034 | 0.9087 | 0.8083 | 0.8712 | 0.8737 | 0.8690 | 0.7972 | 0.8454 |
|  | Area×Circ. | 0.8921 | 0.9081 | 0.9088 | 0.8118 | 0.8688 | 0.8846 | 0.8704 | 0.8004 | 0.8467 |
|  | Minor/Solidity | 0.8863 | 0.9080 | 0.9044 | 0.7959 | 0.8627 | 0.8759 | 0.8666 | 0.7866 | 0.8373 |
|  | MinF/Solidity | 0.8852 | 0.8960 | 0.9062 | 0.8038 | 0.8541 | 0.8584 | 0.8425 | 0.7856 | 0.8498 |
|  | Area×Solidity | 0.8776 | 0.8478 | 0.8884 | 0.8085 | 0.8534 | 0.8222 | 0.8022 | 0.7869 | 0.8194 |
|  | Perim.×Circ. | 0.8902 | 0.9088 | 0.9087 | 0.8108 | 0.8646 | 0.8906 | 0.8702 | 0.8006 | 0.8438 |
|  | A1 (Area×Perim.×Circ.×Solidity×MinF) | 0.8942 | 0.8958 | 0.9074 | 0.8228 | 0.8706 | 0.8690 | 0.8525 | 0.8069 | 0.8515 |
|  | A2 (Area×Perim.×Circ.×Solidity×Minor) | 0.8946 | 0.8984 | 0.9076 | 0.8222 | 0.8714 | 0.8755 | 0.8546 | 0.8078 | 0.8498 |
|  | Kim index | 0.8743 | 0.8390 | 0.8815 | 0.8104 | 0.8489 | 0.8130 | 0.7902 | 0.7810 | 0.8129 |
|  | Mean | 0.8852 | 0.8876 | 0.9013 | 0.8064 | 0.8605 | 0.8597 | 0.8440 | 0.7924 | 0.8373 |
| Enhanced resolution | Area | 0.9053 | 0.8688 | 0.8963 | 0.8617 | 0.8812 | 0.8468 | 0.8204 | 0.8163 | 0.8108 |
|  | Minor | 0.9208 | 0.9433 | 0.9257 | 0.8582 | 0.9069 | 0.9077 | 0.9255 | 0.8524 | 0.8434 |
|  | MinFerret | 0.9159 | 0.9367 | 0.9204 | 0.8456 | 0.9032 | 0.8878 | 0.9241 | 0.8416 | 0.8344 |
|  | Area/perim. | 0.9361 | 0.9421 | 0.9314 | 0.8889 | 0.9225 | 0.9115 | 0.9236 | 0.8642 | 0.8588 |
|  | Area×Circ. | 0.9373 | 0.9463 | 0.9334 | 0.8922 | 0.9228 | 0.9194 | 0.9280 | 0.8714 | 0.8615 |
|  | Minor/Solidity | 0.9149 | 0.9389 | 0.9150 | 0.8471 | 0.8991 | 0.8994 | 0.9212 | 0.8287 | 0.8261 |
|  | MinF/Solidity | 0.9088 | 0.9307 | 0.9084 | 0.8328 | 0.8935 | 0.8777 | 0.9171 | 0.8161 | 0.8148 |
|  | Area×Solidity | 0.9110 | 0.8754 | 0.9052 | 0.8674 | 0.8882 | 0.8527 | 0.8290 | 0.8297 | 0.8242 |
|  | Perim.×Circ. | 0.9362 | 0.9485 | 0.9335 | 0.8932 | 0.9198 | 0.9246 | 0.9304 | 0.8782 | 0.8582 |
|  | A1 (Area×Perim.×Circ.×Solidity×MinF) | 0.9323 | 0.9292 | 0.9279 | 0.8853 | 0.9170 | 0.8978 | 0.9057 | 0.8622 | 0.8568 |
|  | A2 (Area×Perim.×Circ.×Solidity×Minor) | 0.9353 | 0.9338 | 0.9310 | 0.8910 | 0.9201 | 0.9058 | 0.9106 | 0.8691 | 0.8614 |
|  | Kim index | 0.9055 | 0.8690 | 0.8969 | 0.8618 | 0.8819 | 0.8471 | 0.8208 | 0.8159 | 0.8129 |
|  | Mean | 0.9216 | 0.9219 | 0.9188 | 0.8688 | 0.9047 | 0.8898 | 0.8964 | 0.8455 | 0.8386 |

WI and DI: well- and deficit-irrigated, respectively.

Analysis of variance (ANOVA; data not shown) also indicated that the effects of year, mixture treatments, and water stress were very significant on the values of MGW, as well as the two control and 10 selected indices (the total 12 evaluated indices; P<0.0001). As it was expected according to the high correlations between MGW and the image-derived indices, the variation of the indices followed completely the changes in MGW; i.e. the post-anthesis water stress reduced the values significantly (e.g. MGW reduced from 39.291 mg under well-irrigation to 36.157 mg under deficit-irrigation conditions, averaged between two years). In average, MGW also reduced significantly from 39.264 mg in the 1^st^ season to 36.184 mg in the 2^nd^ season (noteworthy, the effect of season on grain yield and most agronomic features were significant. For more information, see Haghshenas et al., 2021). All of the 12 indices showed a similar trend. As a whole, values of MGW and the correlated visual indices were lower in the higher yielding treatments (or conditions) and vice versa; mainly due to the strong negative relationship between grains m^−2^ and MGW on one hand, and the high correlation between grain yield and grains m^−2^ at the other hand (see Haghshenas et al., 2021). The main implication of this observation for the present study was that the variations of the visual indices were highly consistent with those of MGW; regardless of the sources of variation, i.e. significantly different growing seasons, water stress, or mixture treatments.

Figure 4 represents the performance of the linear models developed using the selected indices for predicting MGW (here the images with enhanced resolution were used). As it was expected based on the previous results, all of the ten linear models predicted MGW with a more accuracy compared with the two control indices (RMSE values ranged between 1.003 to 1.201, for the *Area*×*Circ.* and *MinFeret/Solid.* models, respectively). Results of cross-validation and also model parameters have been shown in Table 3. As expected, root mean square errors of cross-validation, followed the pattern of RMSEs reported earlier, i.e. errors of *Area*×*Circ.* < *Perim.*×*Circ.* < *Area/Perim*. Table 3 also represents the reduction percentages of RMSE due to the enhanced resolution by the factor of 10. As a whole, the effect of resolution enhancement was more considerable on the precision of the indices which were based on shape properties (e.g. the products of *circularity*), rather than the size-based features (*Area,* or *MinFeret*).

**Figure 4.**
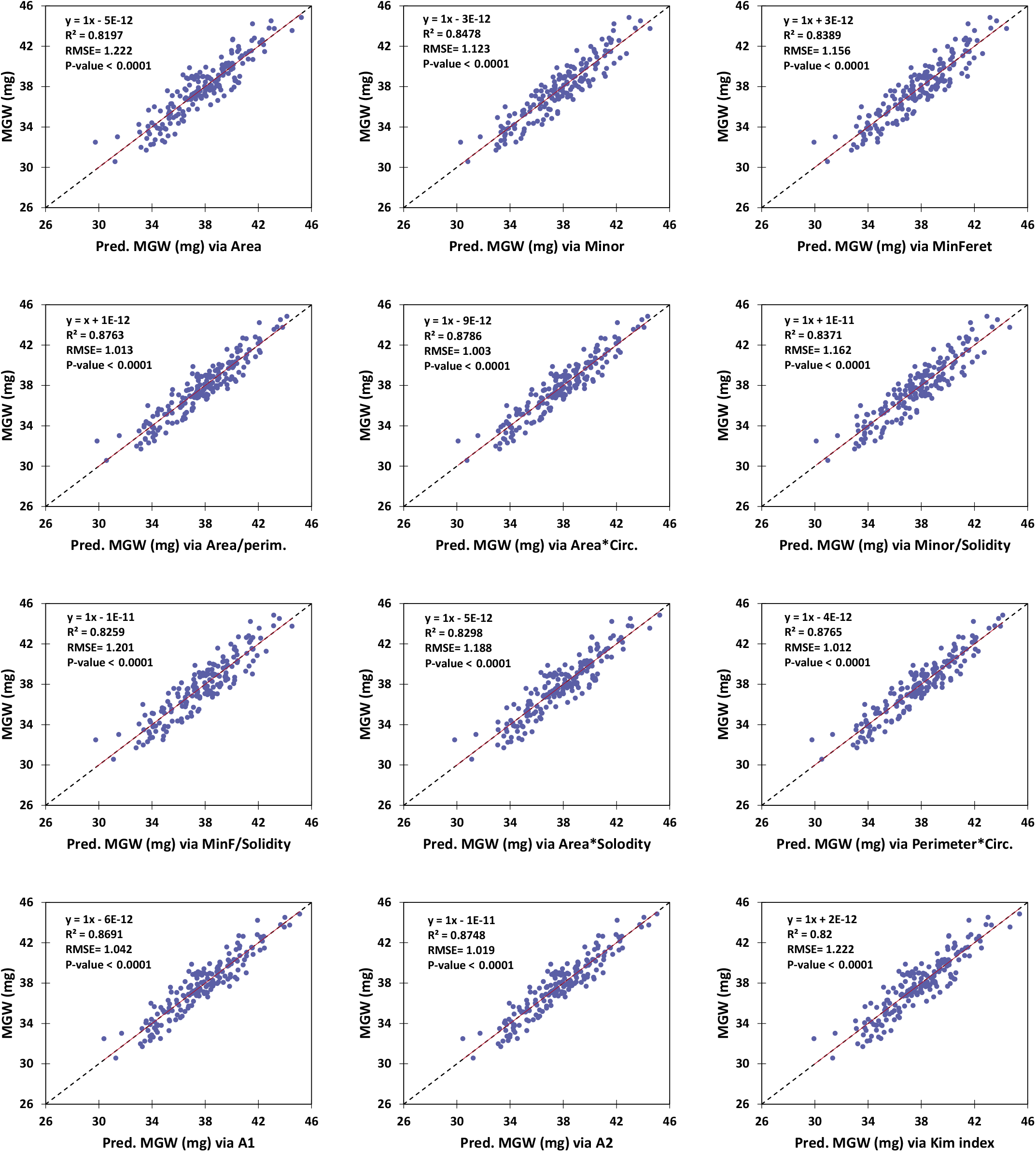
Performance of linear models developed for predicting mean grain weight (MGW) using the superior image-derived indices. The red and dashed lines show the linear trend and 1:1 line, respectively.

**Table 3.**
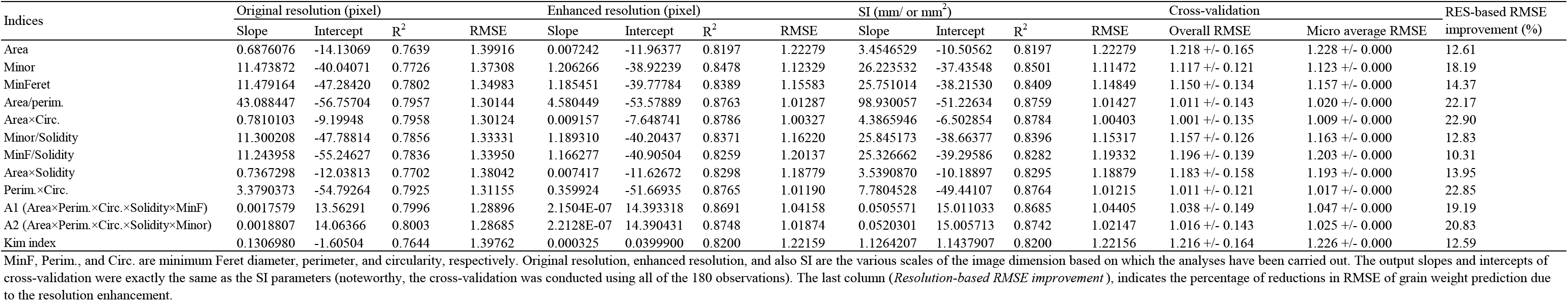
Cross-validation and parameters of the linear models developed for estimation of mean grain weight (MGW; mg) using image-derived indices.

| Indices | Original resolution (pixel) |  |  |  | Enhanced resolution (pixel) |  |  |  | SI (mm/ or mm<sup>2</sup>) |  |  |  | Cross-validation |  | RES-based RMSE improvement (%) |
| --- | --- | --- | --- | --- | --- | --- | --- | --- | --- | --- | --- | --- | --- | --- | --- |
|  | Slope | Intercept | R<sup>2</sup> | RMSE | Slope | Intercept | R<sup>2</sup> | RMSE | Slope | Intercept | R<sup>2</sup> | RMSE | Overall RMSE | Micro average RMSE |  |
| Area | 0.6876076 | -14.13069 | 0.7639 | 1.39916 | 0.007242 | -11.96377 | 0.8197 | 1.22279 | 3.4546529 | -10.50562 | 0.8197 | 1.22279 | 1.218 +/- 0.165 | 1.228 +/- 0.000 | 12.61 |
| Minor | 11.473872 | -40.04071 | 0.7726 | 1.37308 | 1.206266 | -38.92239 | 0.8478 | 1.12329 | 26.223532 | -37.43548 | 0.8501 | 1.11472 | 1.117 +/- 0.121 | 1.123 +/- 0.000 | 18.19 |
| MinFerret | 11.479164 | -47.28420 | 0.7802 | 1.34983 | 1.185451 | -39.77784 | 0.8389 | 1.15583 | 25.751014 | -38.21530 | 0.8409 | 1.14849 | 1.150 +/- 0.134 | 1.157 +/- 0.000 | 14.37 |
| Area/perim. | 43.088447 | -56.75704 | 0.7957 | 1.30144 | 4.580449 | -53.57889 | 0.8763 | 1.01287 | 98.930057 | -51.22634 | 0.8759 | 1.01427 | 1.011 +/- 0.143 | 1.020 +/- 0.000 | 22.17 |
| Area×Circ. | 0.7810103 | -9.19948 | 0.7958 | 1.30124 | 0.009157 | -7.648741 | 0.8786 | 1.00327 | 4.3865946 | -6.502854 | 0.8784 | 1.00403 | 1.001 +/- 0.135 | 1.009 +/- 0.000 | 22.90 |
| Minor/Solidity | 11.300208 | -47.78814 | 0.7856 | 1.33331 | 1.189310 | -40.20437 | 0.8371 | 1.16220 | 25.845173 | -38.66377 | 0.8396 | 1.15317 | 1.157 +/- 0.126 | 1.163 +/- 0.000 | 12.83 |
| MinF/Solidity | 11.243958 | -55.24627 | 0.7836 | 1.33950 | 1.166277 | -40.90504 | 0.8259 | 1.20137 | 25.326662 | -39.29586 | 0.8282 | 1.19332 | 1.196 +/- 0.139 | 1.203 +/- 0.000 | 10.31 |
| Area×Solidity | 0.7367298 | -12.03813 | 0.7702 | 1.38042 | 0.007417 | -11.62672 | 0.8298 | 1.18779 | 3.5390870 | -10.18897 | 0.8295 | 1.18879 | 1.183 +/- 0.158 | 1.193 +/- 0.000 | 13.95 |
| Perim.×Circ. | 3.3790373 | -54.79264 | 0.7925 | 1.31155 | 0.359924 | -51.66935 | 0.8765 | 1.01190 | 7.7804528 | -49.44107 | 0.8764 | 1.01215 | 1.011 +/- 0.121 | 1.017 +/- 0.000 | 22.85 |
| A1 (Area×Perim.×Circ.×Solidity×MinF) | 0.0017579 | 13.56291 | 0.7996 | 1.28896 | 2.1504E-07 | 14.393318 | 0.8691 | 1.04158 | 0.0505571 | 15.011033 | 0.8685 | 1.04405 | 1.038 +/- 0.149 | 1.047 +/- 0.000 | 19.19 |
| A2 (Area×Perim.×Circ.×Solidity×Minor) | 0.0018807 | 14.06366 | 0.8003 | 1.28685 | 2.2128E-07 | 14.390431 | 0.8748 | 1.01874 | 0.0520301 | 15.005713 | 0.8742 | 1.02147 | 1.016 +/- 0.143 | 1.025 +/- 0.000 | 20.83 |
| Kim index | 0.1306980 | -1.60504 | 0.7644 | 1.39762 | 0.000325 | 0.0399900 | 0.8200 | 1.22159 | 1.1264207 | 1.1437907 | 0.8200 | 1.22156 | 1.216 +/- 0.164 | 1.226 +/- 0.000 | 12.59 |

MinF, Perim., and Circ. are minimum Feret diameter, perimeter, and circularity, respectively. Original resolution, enhanced resolution, and also SI are the various scales of the image dimension based on which the analyses have been carried out. The output slopes and intercepts of cross-validation were exactly the same as the SI parameters (noteworthy, the cross-validation was conducted using all of the 180 observations). The last column (*Resolution-based RMSE improvement*), indicates the percentage of reductions in RMSE of grain weight prediction due to the resolution enhancement.

## 4. Discussion

The idea of the present study was exploring more efficient visual indices for wheat MGW, other than 2D grain area. For this purpose, various empirical indices of grain size and shape were evaluated using image processing. It was observed that among the size criteria, one-dimensional indices of grain width (i.e. *Minor* and *MinFeret*) had relatively higher correlations with MGW, compared with the two-dimensional indices of grain area or perimeter (the latter of which was filtered out in the preliminary assessments; R=0.801 when the enhanced-resolution images were used, data not shown). This observation inspired that there might be also other unexplored indices for MGW, which originate from the exclusive physiology of wheat crop, e.g. the processes associated with the grain filling capacity. Therefore, the correlation of MGW with some of the conventional shape indices and also several empirical criteria were tested.

*Area*×*Circ.*, *Perim.*×*Circ.*, and *Area/Perim.* were the superior indices in prediction of MGW using the linear models, and indicated a relatively consistent performance across the various conditions. Furthermore, almost under every of the 4 environmental conditions, other selected indices could predict MGW with a higher precision compared with area. Besides the applicable aspect of this finding, it is also an evidence for the possibility of improving wheat grain weight estimation by exploring new visual indicators.

Based on the formula of the *circularity* index used in ImageJ (see https://imagej.nih.gov/ij/docs/guide/146-30.html), all of the three superior indices have a common factor i.e. the *Area/Perim.* ratio:

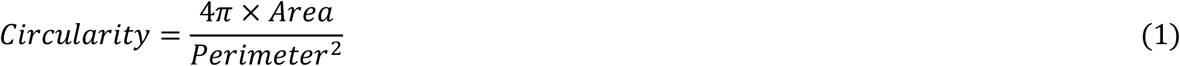

*Therefore*:

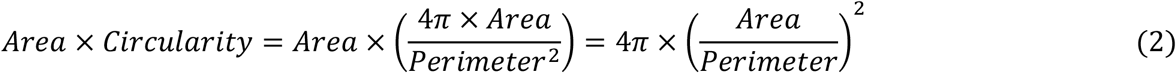

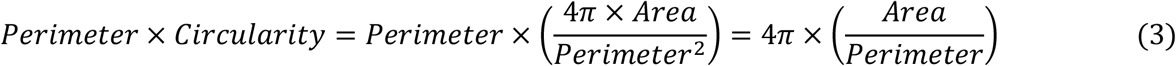

So, the formulae of the two other indices (i.e. *Area*×*Circ.*& *Perim.*×*Circ.*) might be slightly simplified, and consequently the computational cost could be reduced. Such conversions may be particularly important in high-throughput phenotyping; where a considerable number of grains should be analyzed in real-time e.g. using high-speed imaging systems. Besides, these observations imply that the majority of the efficient indices evaluated in the present study are based on two fundamental factors: (i) grain width (measured by *Minor* & *MinFeret*), and (ii) the *Area/Perim.* ratio.

As described before, enhancing the image resolution by the factor of 10 improved the precision of the indices considerably. However, this improvement was not equal for all of the selected indices; as those which were independent of the grain shape, were less influenced (e.g. the size indicators such as *Area* or *MinFeret*, see Table 3). In contrast, the shape-depended indices showed considerably higher degrees of improvement in MGW prediction (for instance, see the indices with the factor of *Circularity*, or even *Minor*, which is resulted from ellipse fitting; see Fig. 1). Therefore, it is necessary to ensure the desirable image resolutions (which is achievable either at the time of imaging/scanning, or using interpolation), before running the analyses.

Noteworthy, since in the present study the analyses were designed and carried out based on the average values, generalization of the findings and models for estimating weight of individual grains might require further assessments. However, considering that each of the 180 samples was consisted of more than 400 grains, it is expected that both types of estimations (MGW and individual grain weight) should be highly correlated. As an evidence for this fact, it was observed that similar to the study of Kim et al. (2021), *Kim index* provided a more precise grain weight estimation than *Area*. More importantly, slopes of the corresponding linear models calculated in both studies were almost similar (see Table 3); despite the differences in the genotypes, treatments, imaging systems, lighting, and probably the image processing algorithms:

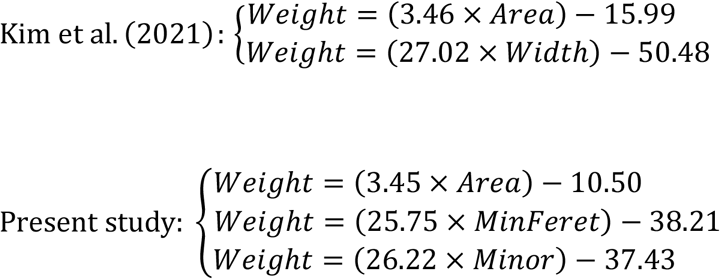

Besides the technical advantageous for developing phenotyping platforms, findings of the present study might also be readily used in wheat physiology and breeding approaches. For instance, the relatively stronger relationship between MGW and grain width (vs. length or even area) may provide valuable implications for the grain filling process; particularly despite the fact that (i) grain filling is an acropetal process and mainly occurs in the grain length direction, and (ii) the 2D grain area provides the information of 2 out of the 3 dimensions. The results also seem to be consistent with the findings of Gegas et al. (2010) who provided the genetic evidences for an emerging phenotypic model where wheat domestication has transformed a long thin primitive grain to a wider and shorter modern grain. Moreover, the superior visual indices introduced in the present study might be used as the selection criteria in breeding programs (e.g. see Alemu et al., 2020); before which the efficiency and stability of the indices should be tested using a more heterogeneous collection of genotypes grown under a broader environmental conditions.

## 5. Conclusion

The present study was conducted to explore the potentially more efficient image-derived indices for predicting MGW of wheat. For this purpose, simple size and shape indices of cultivar mixtures grown under 4 environmental conditions (2 seasons × 2 water conditions) were analyzed. It was observed that MGW had a higher correlation with 10 out of the more than 30 evaluated empirical indices, compared with the well-assessed indicators of projected area (i.e. *Area* & *Kim index*). The best MGW predictions were obtained using the *Area*×*Circ.*, *Perim.*×*Circ.*, and *Area/Perimeter* indices. In general, two main common factors were detected in the majority of the superior indices, i.e. either grain width (evidenced as *Minor* & *MinFeret*) or the *Area/Perimeter* ratio (observed in the simplified forms of *Area*×*Circ.* & *Perim.*×*Circ.* indices). The comparative precision of the ten selected indices was stable under different environmental conditions. Moreover, it was observed that enhancing the image resolution by the factor of 10 could considerably improve the MGW predictions; particularly when the shape-based indices were used. In conclusion, it is expected that the simple predictive linear models developed and validated using the new image-derived indices, could increase the precision of MGW estimations, and also facilitate wheat physiological assessments.

## Acknowledgements

The authors wish to thank Shiraz University for providing field experiment facilities.

## Funding

This research did not receive any specific grant from funding agencies in the public, commercial, or not-for-profit sectors.

## CRediT authorship contribution statement

Abbas Haghshenas: Conceptualization, Methodology, Field experiment, Imaging, Statistical analysis, Visualization, Writing. Yahya Emam: Supervision and reviewing. Saeid Jafarizadeh: Code Writing.

## Declarations of interest

none

## Notes

### Competing Interest Statement

The authors have declared no competing interest.

### Summary of Updates

In the previous version, a key reference (i.e. Kim et al., 2021) was missed.

